# Commensal *Streptococcus mitis* produces two different lipoteichoic acids of type I and type IV

**DOI:** 10.1101/2021.05.06.441962

**Authors:** Nicolas Gisch, Katharina Peters, Simone Thomsen, Waldemar Vollmer, Dominik Schwudke, Dalia Denapaite

## Abstract

The opportunistic pathogen *Streptococcus mitis* possesses, like other members of the Mitis group of viridans streptococci, phosphorylcholine (*P*-Cho)-containing teichoic acids (TAs) in its cell wall. Bioinformatic analyses predicted the presence of TAs that are almost identical with those identified in the pathogen *S. pneumoniae*, but a detailed analysis of *S. mitis* lipoteichoic acid (LTA) was not performed to date. Here we determined the structures of LTA from two *S. mitis* strains, the high-level beta-lactam and multiple antibiotic resistant strain B6 and the penicillin-sensitive strain NCTC10712. In agreement with bioinformatic predictions we found that the structure of one LTA (type IV) was like pneumococcal LTA, except the exchange of a glucose moiety with a galactose within the repeating units. Further genome comparisons suggested that the majority of *S. mitis* strains should contain the same type IV LTA as *S. pneumoniae*, providing a more complete understanding of the biosynthesis of these *P*-Cho-containing TAs in members of the Mitis group of streptococci. Remarkably, we observed besides type IV LTA an additional polymer belonging to LTA type I in both investigated *S. mitis* strains. This LTA consists of β-galactofuranosyl-(1,3)-diacylglycerol as glycolipid anchor and a poly-glycerol-phosphate chain at the *O*-6 position of the furanosidic galactose. Hence, these bacteria are capable of synthesizing two different LTA polymers, most likely produced by distinct biosynthesis pathways. Our bioinformatics analysis revealed the prevalence of the LTA synthase LtaS, most probably responsible for the second LTA version (type I), amongst *S. mitis* and *S. pseudopneumoniae* strains.

## INTRODUCTION

*Streptococcus mitis* is a member of the Mitis group of viridans streptococci (1,2) and is closely related to *S. pneumoniae, S. pseudopneumoniae, S. oralis* and *S. infantis* species (3). The primary habitat of these species is the human upper respiratory tract. *S. mitis* is rarely associated with diseases and presents mostly a non-virulent behavior (3,4). Nevertheless, it can become an opportunistic pathogen via entering the bloodstream and causing bacteremia and/or infective endocarditis (5-7). In contrast, *S. pneumoniae*, which is the closest relative of *S. mitis*, is a well-known respiratory pathogen that is capable of causing a broad spectrum of diseases such as otitis media, bacteremia, pneumonia, and meningitis in humans (8,9). Analyses of the evolution of members of the Mitis group based on genome comparison suggested that *S. mitis, S. pseudopneumoniae* and *S. pneumoniae* arose from a common ancestor, which was a pneumococcus-like organism (10). Both species evolved in parallel either with adaptation to commensal lifestyle (*S. mitis*) by losing virulence-associated genes and genome reduction, or by improving the genomic plasticity necessary for a pathogenic lifestyle (*S. pneumoniae*) (3,11). The species *S. pseudopneumoniae* appears to be genetically intermediary between these two species and was first described in 2004 (11,12).

Lipoteichoic acids (LTA) are main constituents of the Gram-positive cell wall. Their precise role in bacterial physiology has not been fully elucidated, but some LTA-deficient mutants display attenuated virulence in mouse models, defects in biofilm formation or cell division (13,14). In general, LTA contains a lipophilic anchor formed by diacyl-glycerol (DAG), which anchors these molecules to the cell membrane. At the *O*-3 position, DAG carries a glycosyl moiety with attached complex backbone structures consisting of repetitive units (RUs), which are highly variable between different Gram-positive bacteria. Five general LTA types are currently known and are mainly characterized by the architecture of their RUs: polyglycerol-phosphate (type I), complex glycosyl-glycerol-phosphate (type II + III), glycosyl-ribitol-phosphate (type IV), or glycosyl-phosphate (type V) (13). Recently, we showed that the pig pathogen *S. suis* expresses typical type I LTA molecules and complex, mixed-type LTAs, which contain a combination of RUs present in type I and type II/III LTA (15). Type I LTA, the most common and best characterized polymer, has a simple structure consisting of a poly-glycerol-phosphate (poly-Gro-P) chain anchored to a glycolipid in the membrane (13). This type of LTA is found in a many *Firmicutes*, such as *Bacillus subtilis, Staphylococcus aureus, Listeria monocytogenes* and *S. agalactiae* (13,16). Type I LTA was also identified in a *Streptococcus* sp. strain DSM 8747 with the rare glycolipid anchor 3-*O*-(β-D-galactofuranosyl)-1,2-diacylglycerol. DSM 8747 is described as closely related to *S. pneumoniae* (17) although its taxonomic characterization is unpublished and thus the genetic distance to *S. pneumoniae* remained unclear. *S. pneumoniae* produces structurally unique and very complex type IV LTA (18-20). Similar structures are only known for other members of Mitis group of streptococci, such as *S. mitis* SK137 (structure of WTA has been described (21)) and *S. oralis* Uo5 (22). Unlike other bacteria, the TAs in these bacteria and potentially additional members of the Mitis group (such as *S. pseudopneumoniae* and *S. infantis*) share a common biosynthetic pathway as predicted by bioinformatics (23) showing identical chains of both TA polymers (14,19). Remarkably,

*S. mitis* B6 also has a gene encoding a homolog of the LTA synthase LtaS (23). LtaS mediates the polymerization of GroP chains from the head group of the membrane lipid phosphatidylglycerol onto the glycolipid anchor in LTA type I biosynthesis (24,25). The number of LtaS-type enzymes involved in type I LTA synthesis differs between bacterial species. In *S. aureus* LtaS initiates and extends the GroP chain, whereas in *L. monocytogenes* the LTA primase LtaP links the first GroP to the glycolipid anchor, after which LtaS extends the chain (13). This raises the question whether *S. mitis* can produce both type I and type IV LTA. However, up to now the *S. mitis* LTA has not been analyzed.

Here, we report the comprehensive structural analysis of LTA isolated from two *S. mitis* strains, the high-level beta-lactam and multiple antibiotic resistant strain B6 (26) and the penicillin-sensitive strain NCTC10712 (27,28). We determined the LTA structures by chemical degradation, high-resolution mass spectrometry, and one- and two-dimensional, homo- and heteronuclear NMR spectroscopy. We also isolated and analyzed the glycolipids of *S. mitis* B6, verifying the presence of three different glycosylglycerolipids. Furthermore, we assessed key TA biosynthesis genes in the genomes of selected members of the Mitis group of streptococci.

## RESULTS

### Isolation and structural analysis of LTA from *S. mitis* B6 and NCTC10712

LTA from the two *S. mitis* strains B6 and NCTC10712 were prepared according to our previously published workflow for structural analysis (14). LTA shows a tendency to form large aggregates and micelles in water (29) and we showed for *S. pneumoniae* LTA that the first sugar moieties bound to the diacyl-glycerol are not detectable in NMR spectra (18). Therefore, we used *O*-deacylated (hydrazine-treated) and non-aggregating LTA molecules for our analysis. Fig. 1 shows the comparison of ^1^H NMR spectra of *O*-deacylated LTA from the two *S. mitis* strains and *S. pneumoniae*. Spectra have been recorded in deuterated 25 mM sodium phosphate buffer (pH 5.5) at 300 K. The adjustment of the pH value of the NMR solvent was necessary for a homogeneous degree of protonation of the amino group at position 4 of the 2-acetamido-4-amino-2,4,6-trideoxygalactose (AATGal*p*), making the NMR chemical shifts better comparable and reproducible between multiple samples. The spectra of *O*-deacylated LTA from the two *S. mitis* strains were almost identical but slightly different to the spectrum of *O*-deacylated LTA from *S. pneumoniae* strain D39Δ*cps*Δ*lgt* (this strain lacks the capsular polysaccharide and the gene encoding for the lipoprotein diacylglyceryl transferase (Lgt) and is therefore deficient in lipidation of pre-lipoproteins; its LTA has been shown to be identical with that of the D39 wild type (14,18)), indicating different composition in type IV LTA (see dashed lines in magnification of the anomeric region). Interestingly, the *S. mitis* spectra also showed the presence of an additional polymer that was absent in *S. pneumoniae* and is represented, e.g., by an additional anomeric proton at δ_H_ 5.02 ppm (black arrows, Fig. 1) and intensive signals between δ_H_ 4.10-3.85 ppm (marked with *, Fig. 1). Further NMR analyses then proved the presence of a second type of LTA. In addition to the ribitol-phosphate containing LTA known from pneumococci, the NMR signals identified a glycerol-phosphate containing polymer. The best approach to compare these classes is by their ^31^P NMR spectra (Fig. 2). The broad signal in the ^31^P NMR spectra from 1.48-1.12 ppm (B6) or 1.39-1.10 ppm (NCTC10712) was only present in *O*-deacylated LTA from *S. mitis*, coming from poly-glycerol-phosphate chains. The shift of signals for the phosphates in ribitol-5-*P* moieties from 1.89/1.80 ppm in pneumococcal *O*-deacylated LTA to 1.64/1.52 ppm (B6) and 1.61/1.48 ppm (NCTC10712) in *S. mitis* further points to a deviating carbohydrate content in the type IV LTA. The same chemical shifts for the phosphates of the *P*-Cho moieties reflect the identity of the structure in this part of the molecules. Fig. 3 shows the ^1^H, ^13^C-heteronuclear single quantum correlation (HSQC) NMR of LTA_N2H4_ of *S. mitis* strain B6 and the signal assignment as an example for both strains, since almost identical spectra were obtained. The complete NMR chemical shift data for type I and type IV *O*-deacylated *S. mitis* LTA are summarized in Tables 1 and 2. The *O*-deacylated lipid anchor of the type IV LTA is α-glucopyranosyl-glycerol. The following RUs all have the pseudo-pentasaccharide composition (→4)-6-*O*-*P*-Cho-α-D-Gal*p*NAc-(1→3)-6-*O*-*P*-Cho-β-D-Gal*p*NAc-(1→1)-Rib-ol-5-*P*-(*O*→6)-β-D-Gal*p*-(1→3)-AATGal*p*(1→). The first RU (RU 1) attached to the lipid anchor is β-linked, whereas all other RUs, representing an elongation of the carbohydrate part in the LTA molecule, are α-1-linked to the precedent RU. The *O*-deacylated lipid anchor of the type I LTA is β-galactofuranosyl-glycerol, which is elongated by the poly-glycerol-phosphate chain at the *O*-6 position of the Gal_*f*_.

**Figure 1.**
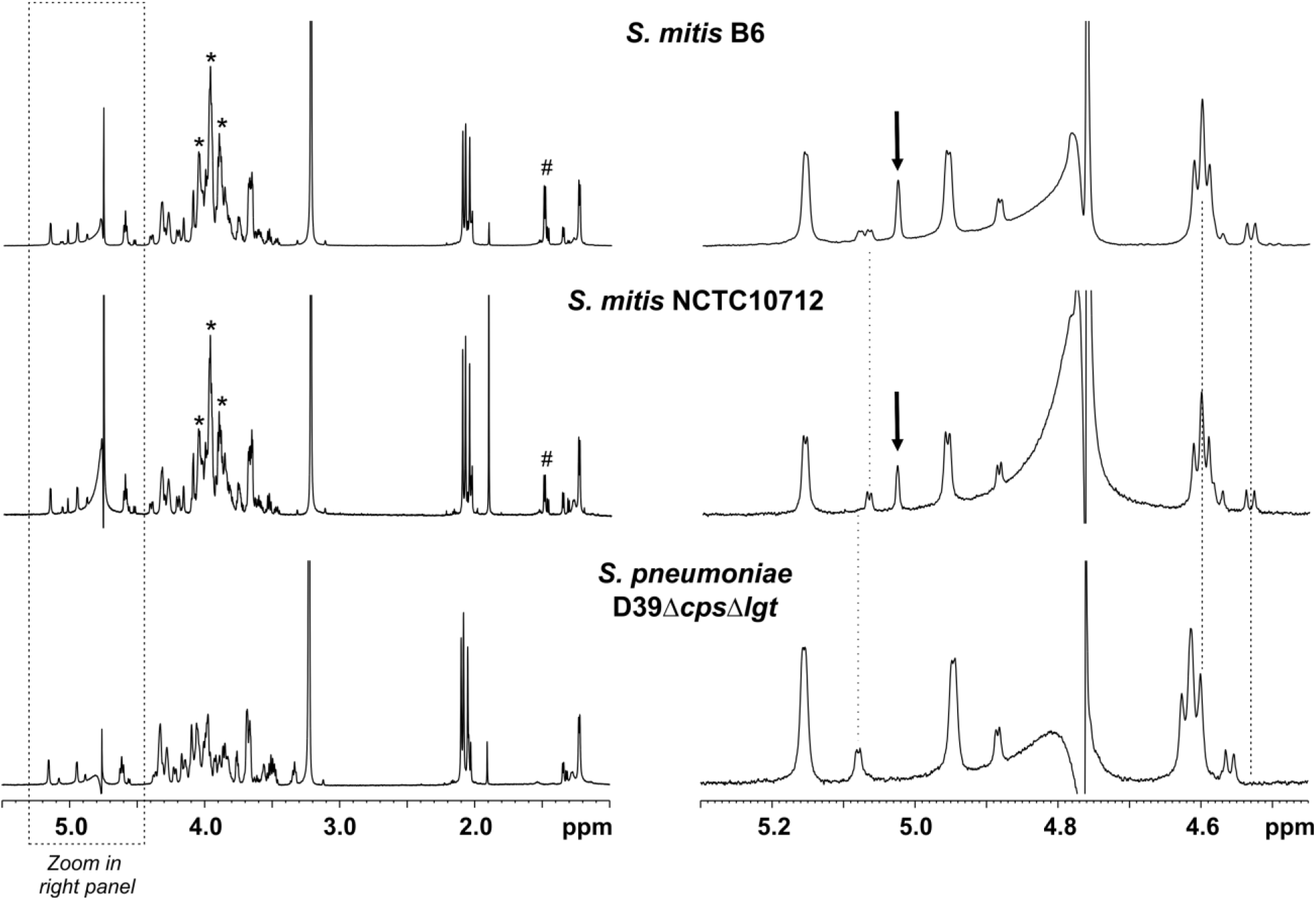
Identification of two LTA polymers in *S. mitis* strains. Shown are ^1^H NMR spectra (δ_H_ 5.5-1.0 in left panel; (δ_H_ 5.3-4.45 (anomeric region) in right panel) of *O*-deacylated LTA of *S. mitis* strains B6 (top) and NCTC10712 (middle) as well as from *S. pneumoniae* D39Δ*cps*Δ*lgt* (bottom), all recorded in deuterated 25 mM sodium phosphate buffer (pH 5.5) at 300 K. Dotted lines in right panel indicate deviating anomeric signals in the different LTA preparations. Stars in left panel as well as black arrows in right panel indicate the presence of an additional polymer in the *S. mitis* LTA preparations, which is absent in *S. pneumoniae*. Signals marked with # (right panel) present in *S. mitis* LTA preparations result from a molecule that is formed during the hydrazine-mediated alanine-cleavage (most likely alanine-hydrazide), which was removed in the *S. pneumoniae* sample by chromatography (18).

**Figure 2.**
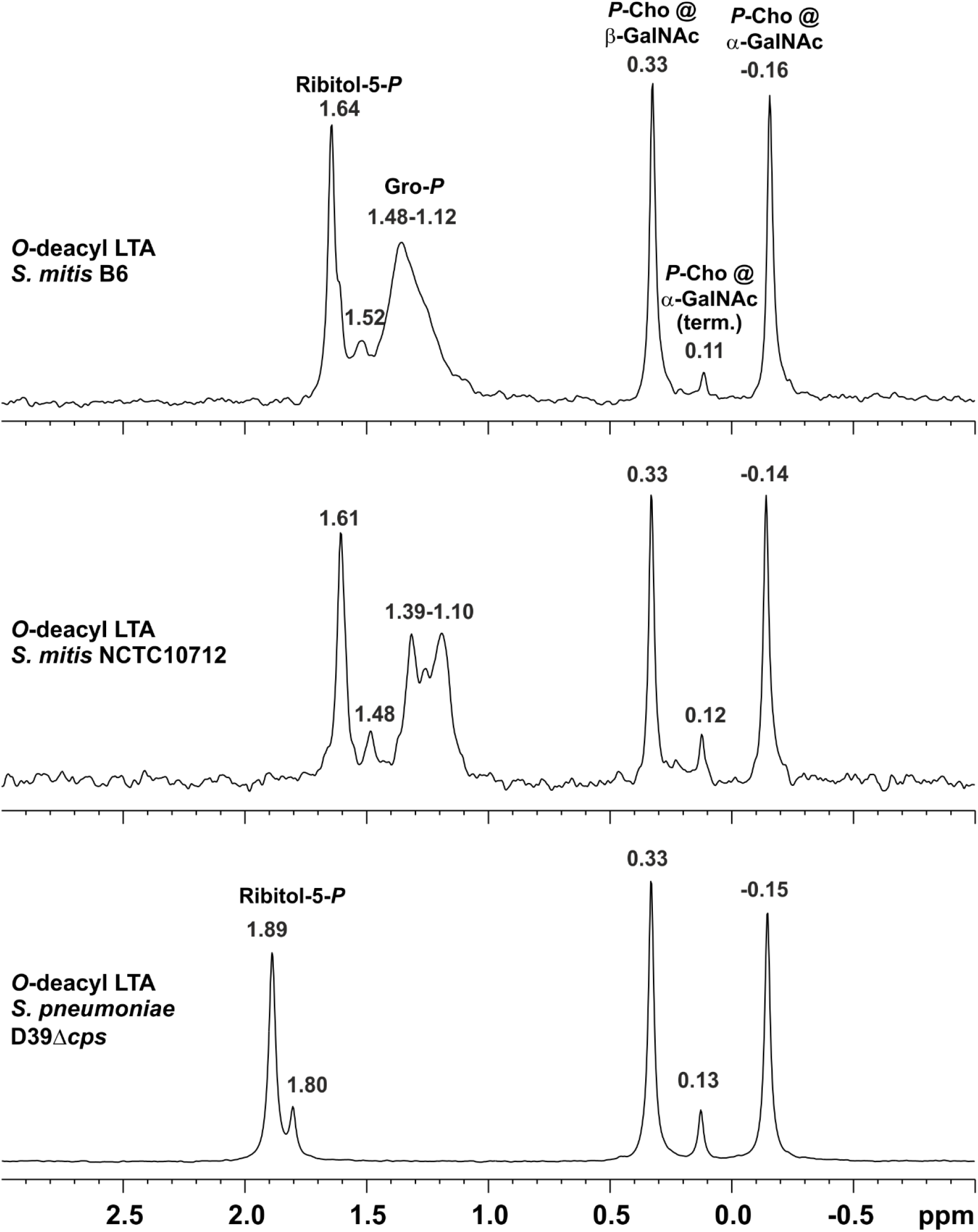
Presence of a poly-glycerolphosphate containing LTA in *S. mitis*. Shown are ^31^P NMR spectra (δ_P_ 3.0-(−1.0)) of *O*-deacylated LTA of *S. mitis* strains B6 (top) and NCTC10712 (middle) as well as from *S. pneumoniae* D39Δ*cps* (bottom), recorded in D_2_O at 300 K. The broad signal between δ_P_ 1.48 and 1.12 ppm (B6) or 1.39 and 1.10 ppm (NCTC10712), respectively, only present in *O*-deacylated LTA from *S. mitis*, clearly points to the presence of poly-glycerol-phosphate chains (compare (15)).

**Figure 3.**
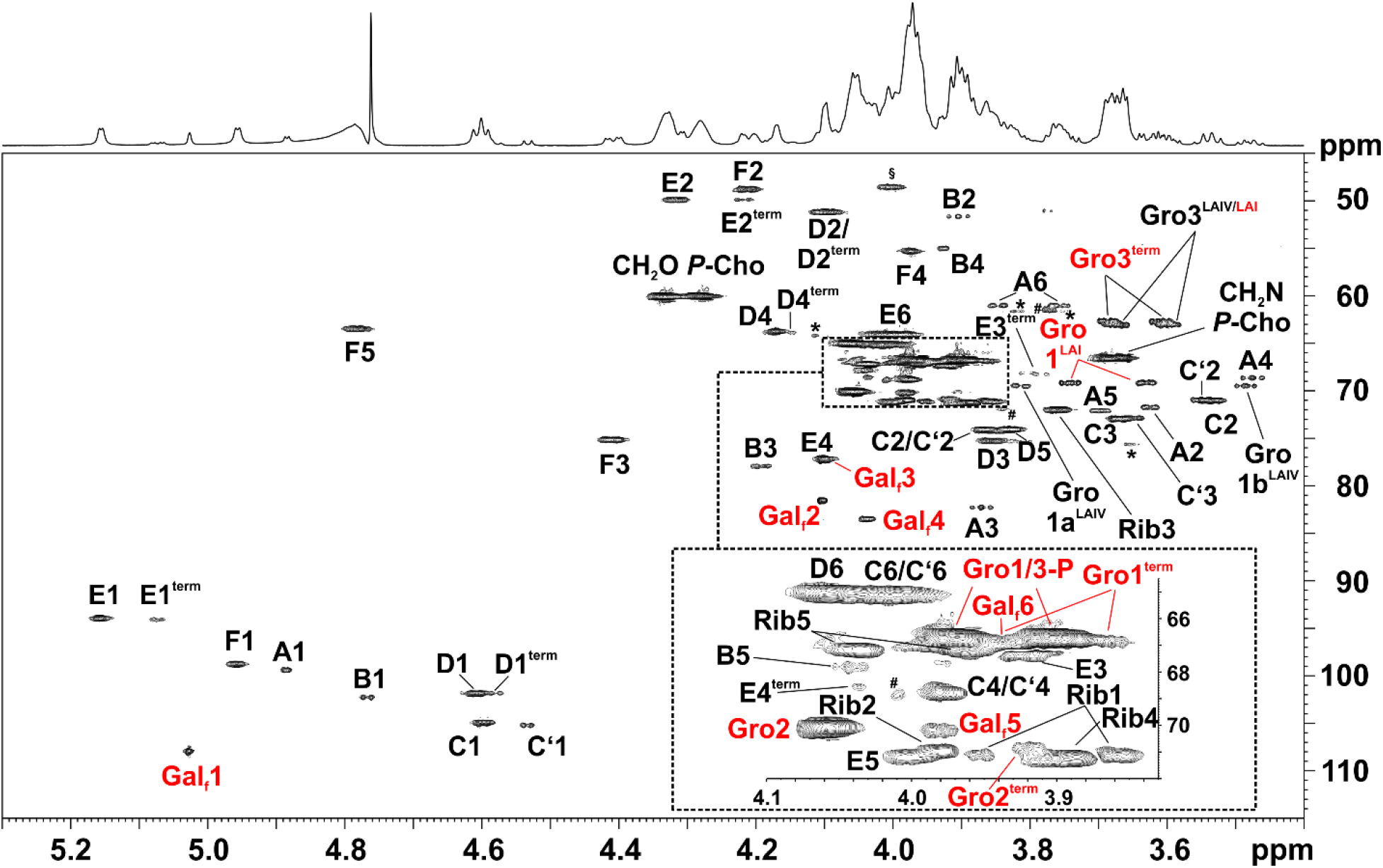
NMR analysis of *O*-deacylated LTA of *S. mitis* strain B6. Shown is a section (δ_H_ 5.30-3.40; δ_C_ 115-45) of the ^1^H,^13^C-HSQC NMR spectrum (recorded in deuterated 25 mM sodium phosphate buffer (pH 5.5) as dept-version) obtained from *O*-deacylated LTA of *S. mitis* strain B6 including assignment of signals (red: type I LTA; black: type IV LTA). The corresponding NMR chemical shift data are listed in Table 1 (type I LTA) and Table 2 (type IV LTA), respectively.

**Table 1.** **^1^H (700.4 MHz), ^13^C NMR (176.1 MHz), and ^31^P NMR (283.5 MHz) chemical shift data (d, ppm) [*J*, Hz] for *S. mitis* strain B6 type I LTA after hydrazine treatment (*O*-deacyl LTA) recorded in deuterated 25 mM sodium phosphate buffer (pH 5.5) at 300 K.**

| Residue<br>(assignment) | H-1<br>*C-1* | H-2<br>*C-2* | H-3<br>*C-3* | H-4<br>*C-4* | H-5<br>*C-5* | H-6<br>*C-6* |
| --- | --- | --- | --- | --- | --- | --- |
| Gro-(1→[Gro<sup>LAI</sup>]) | 3.76-3.73*<br>3.65-3.61*<br>69.2 | 3.97-3.93*<br>*n.d.* | 3.70-3.65*<br>3.62-3.58*<br>63.2 |  |  |  |
| →1)-β-D-Galf-(6→P [Galf] | 5.04-5.02*<br>107.9 | 4.11-4.09*<br>81.6 | 4.10-4.07*<br>77.4 | 4.05-4.01*<br>83.5 | 3.99-3.96*<br>70.2 | 3.96-3.92*<br>67.1 [5.3] |
| P→1)-Gro-(3→P [Gro-P] | 4.00-3.95*<br>3.93-3.87*<br>66.9 [6.4] | 4.08-4.03*<br>70.1 | 4.00-3.95*<br>3.93-3.87*<br>66.9 [6.4] |  |  |  |
| P→1)-Gro [Gro<sup>term</sup>] | 3.95-3.91*<br>3.89-3.85*<br>67.0 [5.2] | 3.93-3.90*<br>70.9 | 3.70-3.66*<br>3.63-3.59*<br>62.7 |  |  |  |
| <sup>31</sup>P# | Gal-6-P-Gro *n.d.*; Gro-P-Gro 1.48-1.12. |  |  |  |  |  |

#shifts from spectra recorded in D<sub>2</sub>O @ 300 K (see Fig. 2); \*non-resolved multiplet; *n.d.* = not detected.

**Table 2.**
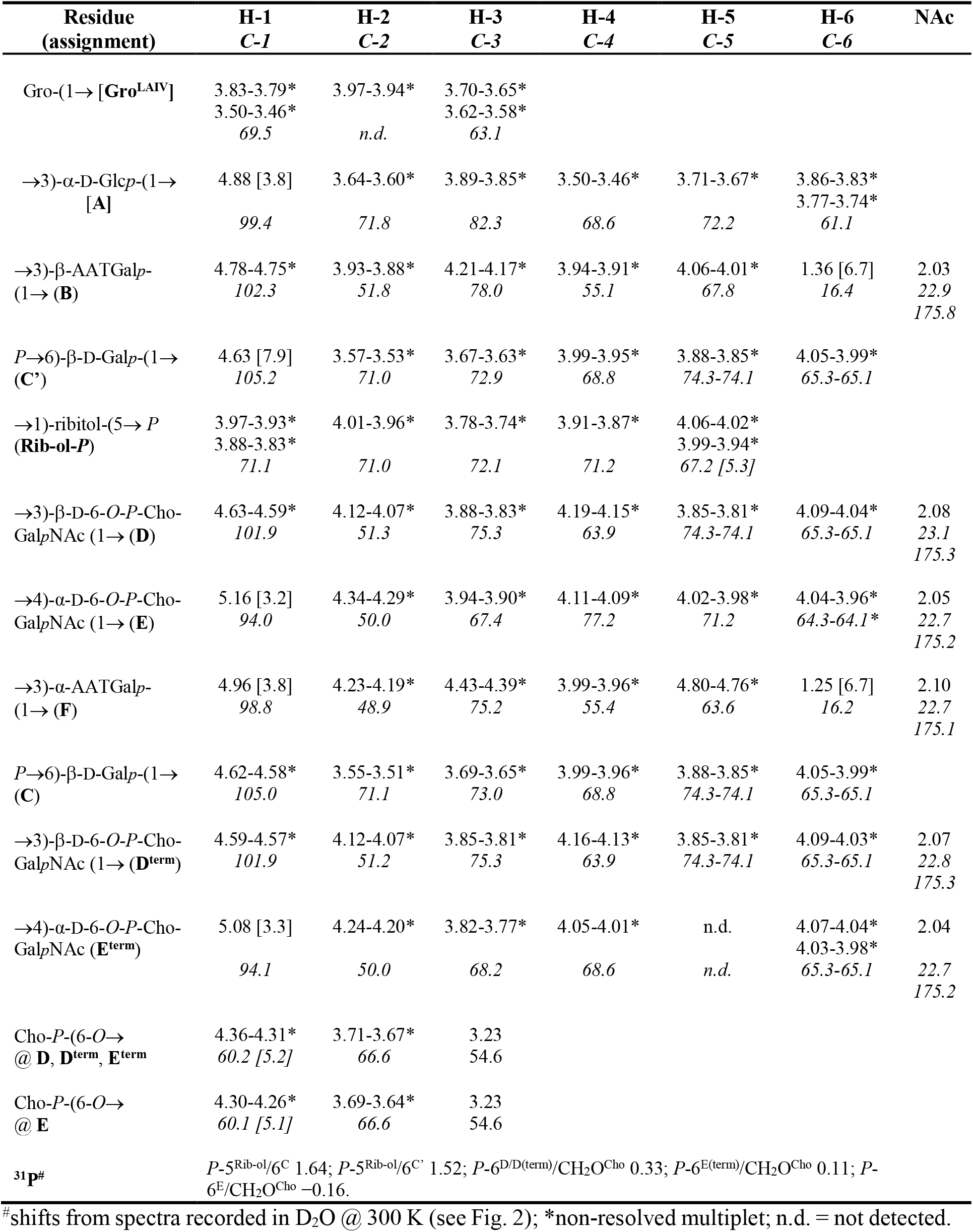
**^1^H (700.4 MHz), ^13^C NMR (176.1 MHz), and ^31^P NMR (283.5 MHz) chemical shift data (d, ppm) [*J*, Hz] for *S. mitis* strain B6 type IV LTA after hydrazine treatment (*O*-deacyl LTA) recorded in deuterated 25 mM sodium phosphate buffer (pH 5.5) at 300 K.**

The two LTA polymers are present in almost equally amounts in both strains. As judged by integration of the ^1^H NMR signals for Gal_*f*_ H-1 (for type I LTA) and C’ H-1 (for type IV LTA) the ratio was appr. 55:45 in both preparations of strain NCTC10712. The ratio in strain B6 was in one preparation 60:40, in the other 40:60. To verify the two different LTA types, we analyzed the *O*-deacylated LTA preparations from the *S. mitis* strains using high-resolution mass spectrometry. Due to very different ionization efficiencies of the two LTA types, we adjusted the recorded mass range in a way that each LTA type was monitored separately. Type I LTA molecules ionized as doubly or triply charged species and were recorded in an *m/z*-window of 500 to 1650. By contrast, type IV LTA molecules ionized as quadruple and quintuple charged species and were recorded in an *m/z*-window of 1600 to 3000. Representative charge deconvoluted spectra are depicted in Fig. 4. The identified molecular species are listed in Tables 3 and 4, respectively. The LTA of strain NCTC10712 lacks the molecular species displaying a complete *P*-Cho substitution pattern, the most abundant molecules present lack one *P*-Cho substituent (Fig. 4B). This is due to a more pronounced *P*-Cho hydrolysis of the terminal RU mediated by the phosphorylcholine esterase Pce, which is also indicated by the shift of the ^1^H NMR signal representing the anomeric proton of the terminal α-D-6-*O*-*P*-Cho-Gal*p*NAc (**E**^**term**^) from δ_H_ 5.08 ppm to δ_H_ 5.07 ppm (Fig. 1, right panel). We recently reported that Pce is the only enzyme that modifies the *P*-Cho substitution pattern of the TAs in *S. pneumoniae* (30). This effect was not observed in a second, independent LTA preparation of this strain. In this second preparation less *P*-Cho hydrolysis and a tendency to longer chain length was observed (Fig. S1). However, these differences did not impact the general structural composition of the LTA. Finally, the ^1^H NMR analysis of the native LTA preparation of *S. mitis* NCTC10712 (Fig. S2) revealed the presence of alanine residues on the LTA molecules, as is observed in *S. pneumoniae*. The chemical structures of the two LTA polymers identified in the two analyzed *S. mitis* strains are shown in Fig. 5.

**Figure 4.**
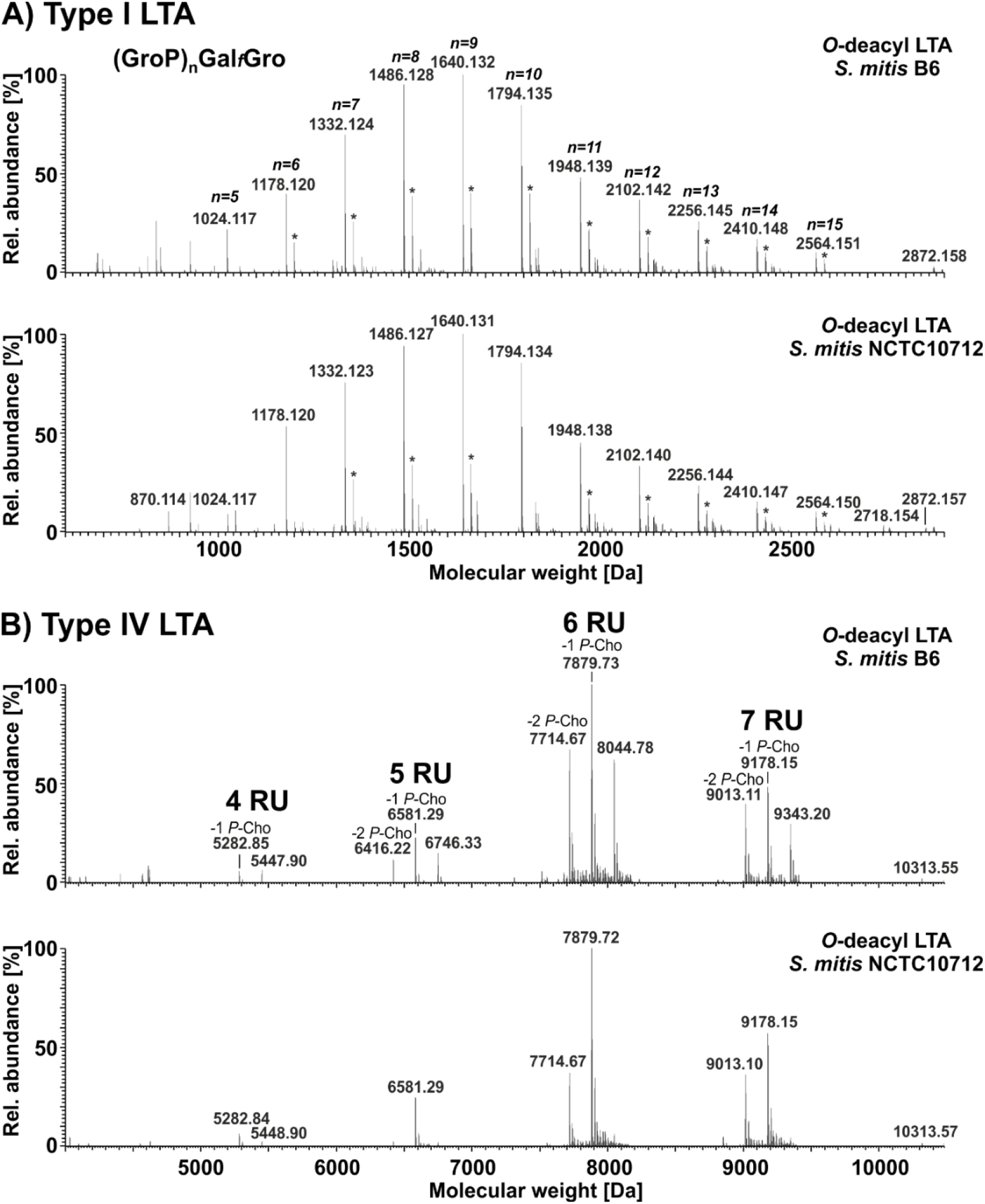
Molecular species distribution of *O*-deacylated LTA of *S. mitis* strains NCTC10712 and B6. **A**) Charge-deconvoluted spectra of a representative MS-analysis for each strain performed in the negative ion mode recorded in an *m/z*-window of 500 to 1650, focusing on the ionization of the type I LTA. Signals marked with * represent major abundant sodium adducts (Δ*m* = +21.98 Da). n = number of GroP repeats. **B)** Charge-deconvoluted spectra of a representative MS-analysis for each strain performed in the negative ion mode recorded in an *m/z*-window of 1600 to 3000, focusing on the ionization of the type IV LTA. Details of all assigned molecular species are summarized in Table 3 (type I LTA) and Table 4 (type IV LTA), respectively. Relative abundance for a spectral region was always normalized to the respective base peak.

**Table 3.**
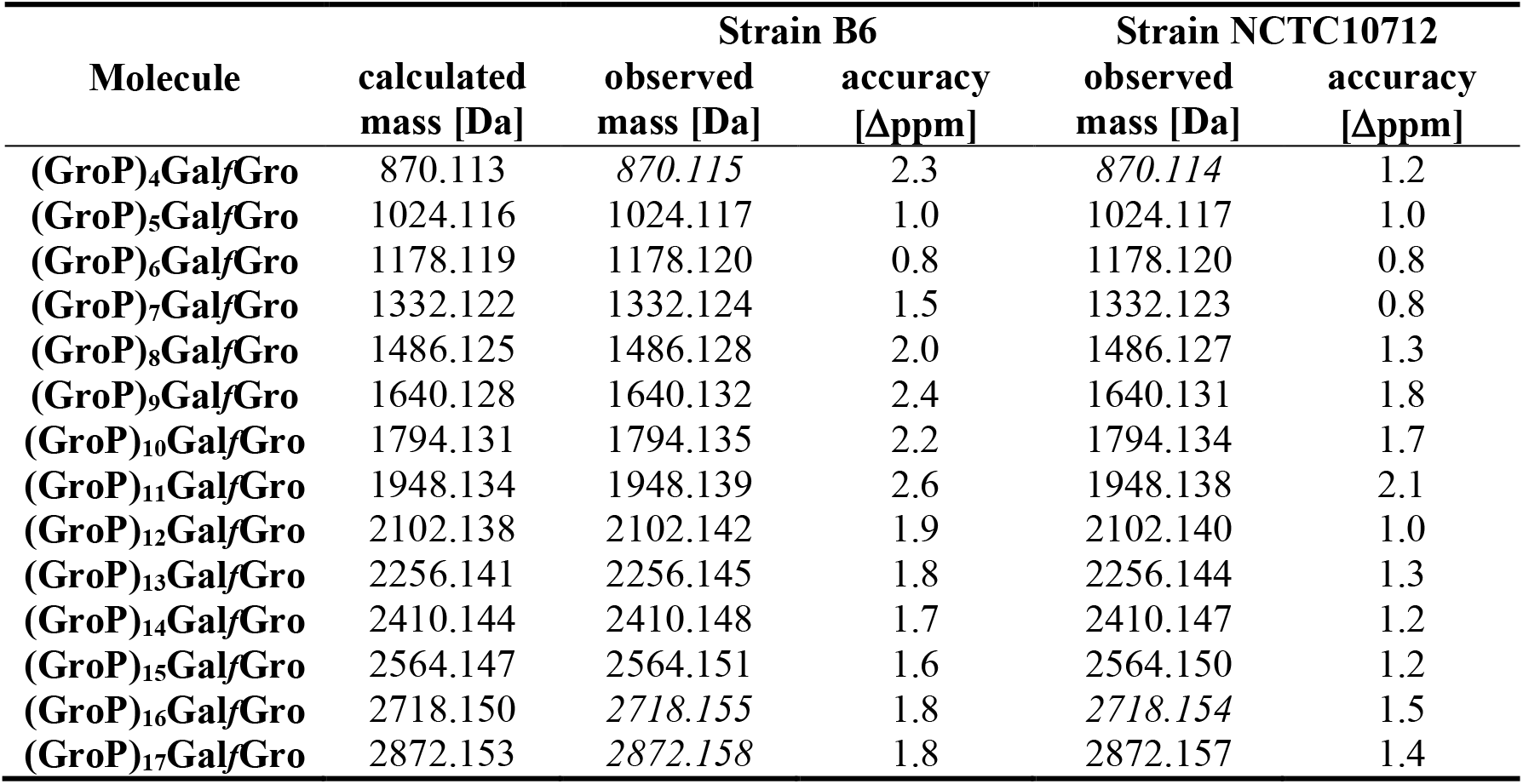
Mass spectrometric analysis of *O*-deacyl type I LTA of *S. mitis* strains B6 and NCTC10712. Summary of calculated monoisotopic neutral masses and observed molecular masses [Da] for LTA preparations after hydrazine treatment. For each preparation, independent MS analyses have been performed and identified molecules are listed as a combined list. Masses observed only in one of the two preparations of a strain are written in italic style. Accuracy of the measurement is stated as Δppm.

**Table 4.** Mass spectrometric analysis of *O*-deacyl type IV LTA of *S. mitis* strains B6 and NCTC10712. Summary of calculated monoisotopic neutral masses and observed molecular masses [Da] for LTA preparations after hydrazine treatment. For each preparation, independent MS analyses have been performed. Identified molecules from one analysis per strain (spectra shown in Fig. 4) are listed. Masses observed only in very minor intensity are written in italic style. Accuracy of the measurement is stated as Δppm; n.d. = not detected; ^#^ = 1^st^ isotopic peak, ^§^ = 2^nd^ isotopic peak.

| Molecule | calculated mass [Da] | Strain B6 |  | Strain NCTC10712 |  |
| --- | --- | --- | --- | --- | --- |
| | | observed mass [Da] | accuracy [Δppm] | observed mass [Da] | accuracy [Δppm] |
| **4 RU** | 5447.89 | 5447.90 | 1.8 | *5448.90*<sup>#</sup> | 1.8 |
| - 1 *P-Cho* | 5282.83 | 5282.85 | 3.8 | 5282.84 | 1.9 |
| - 2 *P-Cho* | 5117.78 | n.d. | - | n.d. | - |
| **5 RU** | 6746.34 | 6746.33 | -1.5 | *6746.36* | 3.0 |
| - 1 *P-Cho* | 6581.28 | 6581.29 | 1.5 | 6581.29 | 1.5 |
| - 2 *P-Cho* | 6416.23 | 6416.22 | -1.6 | *6416.24* | 1.6 |
| **6 RU** | 8044.78 | 8044.78 | 0.0 | *8045.78*<sup>#</sup> | -1.2 |
| - 1 *P-Cho* | 7879.73 | 7879.72 | -1.3 | 7879.72 | -1.3 |
| - 2 *P-Cho* | 7714.67 | 7714.67 | 0.0 | 7714.67 | 0.0 |
| **7 RU** | 9343.23 | 9343.20 | -3.2 | 9343.13 | -10.7 |
| - 1 *P-Cho* | 9178.18 | 9178.15 | -3.3 | 9178.15 | -3.3 |
| - 2 *P-Cho* | 9013.12 | 9013.11 | -1.1 | 9013.10 | -2.2 |
| **8 RU** | 10641.68 | n.d. | - | n.d. | - |
| - 1 *P-Cho* | 10476.62 | n.d. | - | *10477.58*<sup>#</sup> | -4.8 |
| - 2 *P-Cho* | 10311.57 | *10313.55*<sup>§</sup> | 1.9 | *10313.57*<sup>§</sup> | 0.0 |

**Figure 5.**
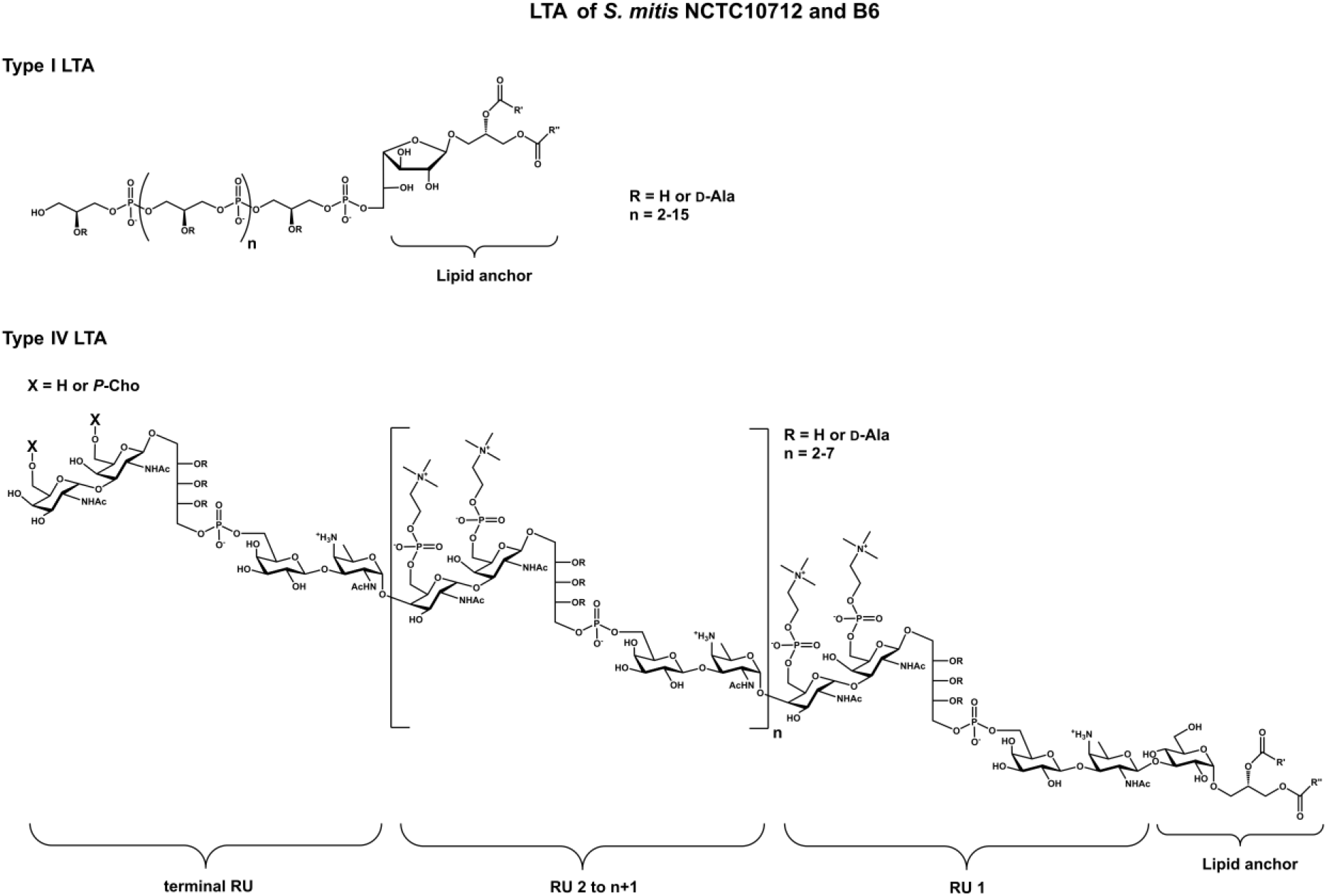
Chemical structures of LTA isolated from *S. mitis* strains NCTC10712 and B6.

### Investigation of the glycosylglycerolipids from *S. mitis* B6

As they represent the putative cell wall anchor motifs for LTA molecules, we also isolated the glycosylglycerolipids from *S. mitis* strain B6 by preparative thin-layer chromatography. The ^1^H NMR spectra of the pool of mono-glycosyl-diacylglycerols and di-glycosyl-diacylglycerols are depicted in Fig. 6, respectively. Our analysis revealed the presence of two different mono-glycosyl-diacylglycerols, α-glucopyranosyl-(1,3)-diacylglycerol and β-galactofuranosyl-(1,3)-diacylglycerol, and a single di-glycosyl-diacylglycerol, α-galactopyranosyl-(1,2)-α-glucopyranosyl-(1,3)-diacylglycerol. The complete NMR chemical shift data for the identified glycosylglycerolipids are summarized in Table 5.

**Figure 6.**
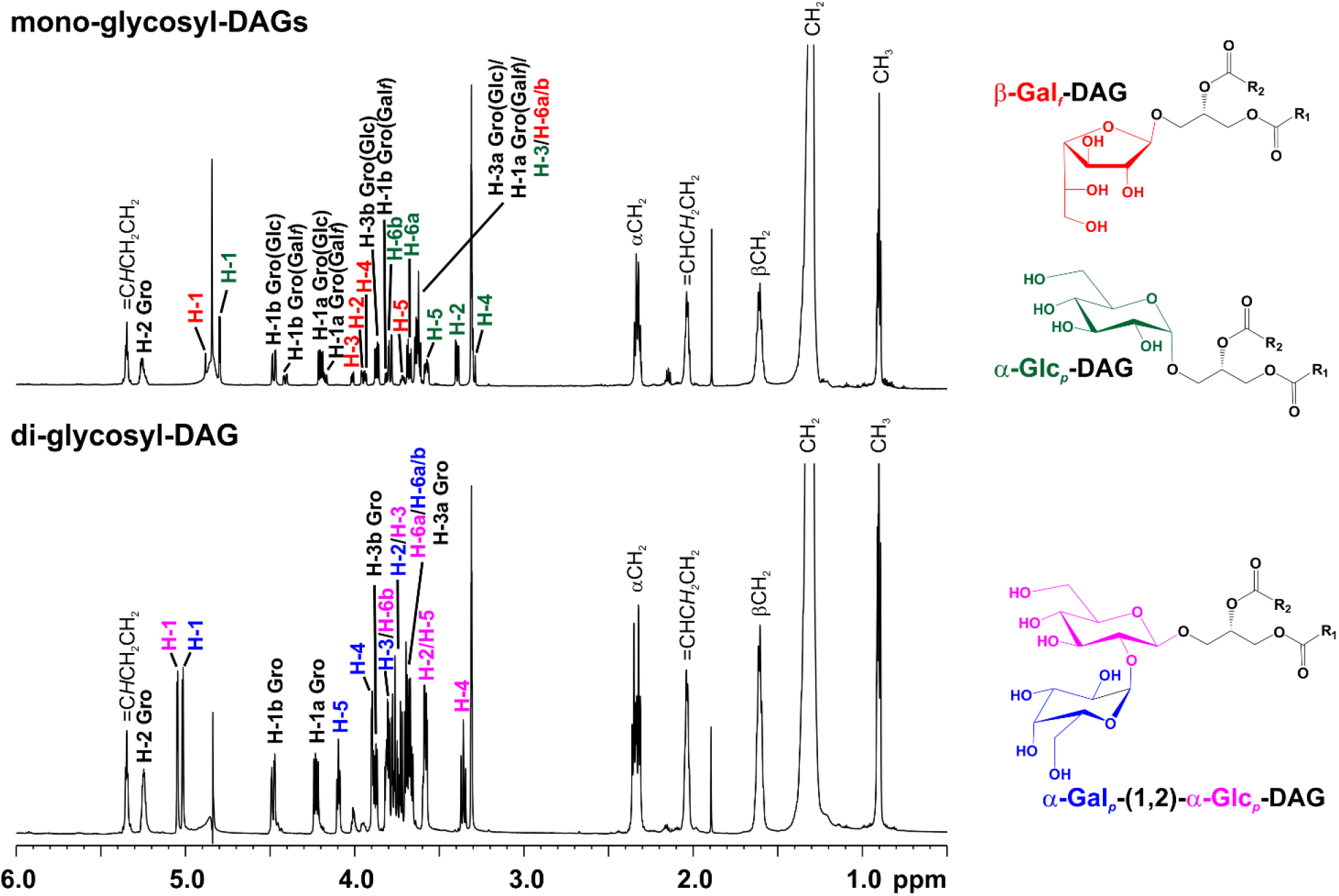
^1^H NMR analysis of glycosylglycerolipids isolated from *S. mitis* strain B6. Shown are ^1^H NMR spectra (δ_H_ 6.0-0.5) of glycosylglycerolipids (*top*: mixture of two mono-glycosyl-diacylglycerols; *bottom*: one di-glycosyl-diacylglycerol) isolated from *S. mitis* strain B6, recorded in MeOH-*d*_*4*_ at 300 K. The corresponding NMR chemical shift data are listed in Table 5, the specific chemical structures of the observed glycosylglycerolipids are depicted right hand in this figure.

**Table 5.**
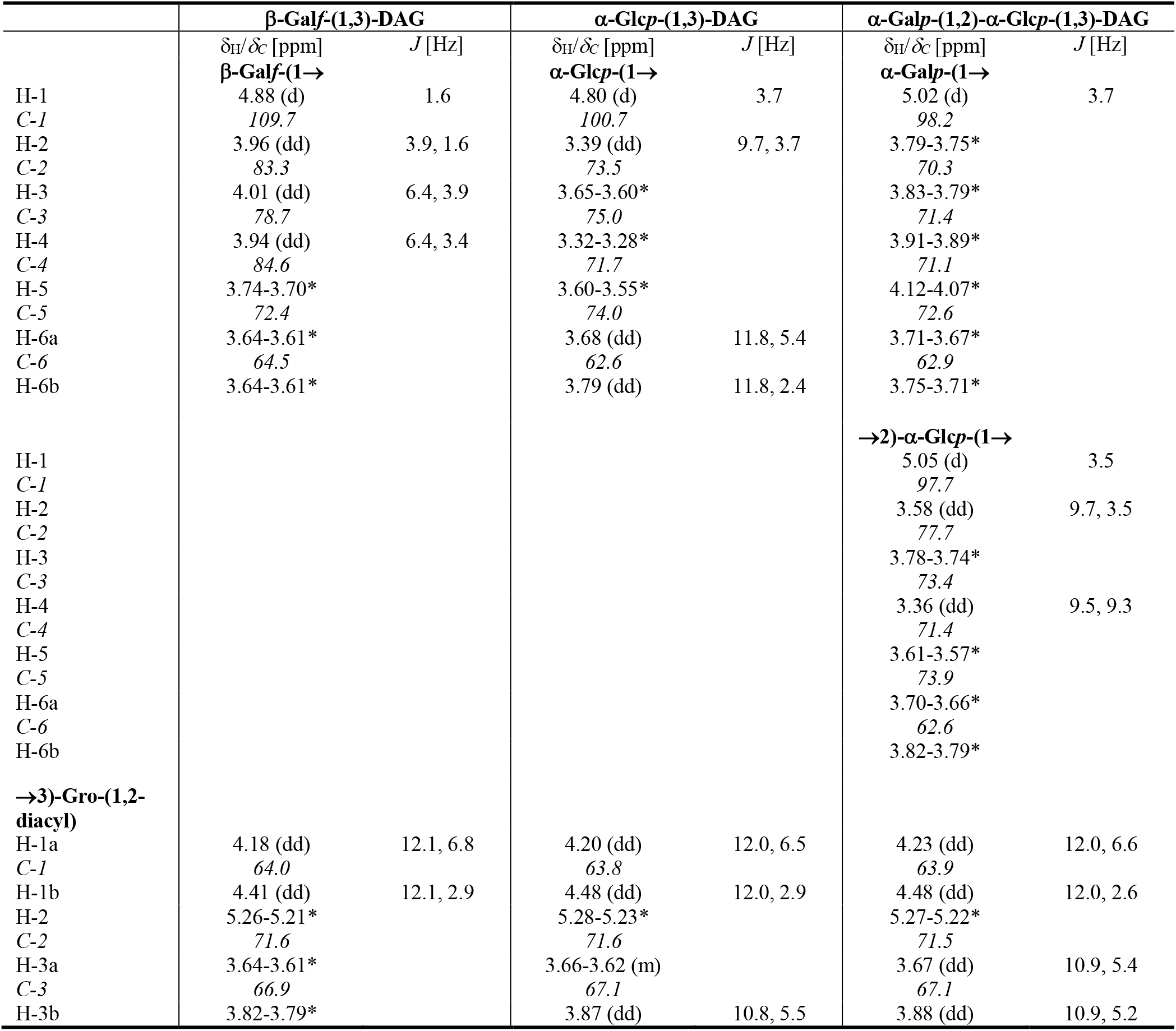
^1^H (700.4 MHz) and ^13^C NMR (176.1 MHz) chemical shift data (d, ppm) [*J*, Hz] for the carbohydrate moieties of glycosylglycerolipids isolated from *S. mitis* strain B6 recorded in MeOH-*d*_*4*_ at 300 K. *non-resolved multiplet.

### *In silico* analysis of LTA biosynthesis key enzymes in *S. mitis* and closely related species

The TAs in members of the Mitis group like *S. mitis, S. pneumoniae* and *S. oralis* share a common biosynthetic pathway (23), which therefore have structurally identical carbohydrate chains in both of their TAs – LTA and WTA – and the analysis of one of the polymers can be considered as representative for both. For *S. mitis* only the structural analysis of the WTA of strain SK137 has been described. In this strain a β-D-glucose has been identified within the RU (21), which was identical to that in *S. pneumoniae* (14,18). Our structural analysis of the LTA of *S. mitis* strains B6 and NCTC10712 revealed the presence of a β-D-galactose at this position, which was in line with the expectation based on the gene content of these strains. Hence, we raised the question about the predominant structural type in *S. mitis* strains. BLAST searches in the NCBI microbial genomes database revealed that 68.8% of the *S. mitis* strains (95 out of 138 strains) possessed the equivalent of the *S. mitis* B6 *smi_1983* gene, which shows 94% homology to *sp70585_0164* of *S. pneumoniae* 70585 (Table S1.1). The *S. mitis* type strain NCTC12261 (also named ATCC 49456) is identical to most *S. mitis* strains in this respect. 36 strains (26.8%) were identical to Spr0091/SPD_0098 of *S. pneumoniae* R6/D39 and therefore likely incorporate a glucose into their TA. *S. mitis* SK137 strain shows homology to Spr0091 (97% identity), which is consistent with the reported structure (21). Only seven *S. mitis* strains (5.1%) did not reveal an identical gene, which is probably due to incompleted genome sequences or a potential mislabeling of these strains. Indeed six out of seven of these genomes do not contain any genes encoding proteins involved in the biosynthesis of type IV LTA (Table S1.1), hence these were not further considered in this study.

In order to ascertain the homogeneity of the *S. mitis* group, we focused for further analysis on four different enzymes, two from *S. mitis* B6 and two from *S. oralis* Uo5, which are essential for the biosynthesis of the respective type IV TA and searched for homologs by tBlastn analysis. The selected enzymes were the teichoic acid flippase TacF (Smi_1229; Spr1150 in *S. pneumoniae* R6 (23)) and the repeating unit (RU) polymerase (Smi_0770; Spr1222 in *S. pneumoniae* R6 (23)) for *S. mitis*. As determinants for an *S. oralis*-like TA structure we chose the *S. oralis* Uo5 enzymes TacF (Sor_0765) and LicD4 (Sor_0762). The latter has no homolog encoded from the *S. mitis*/*S. pneumoniae* genomes. LicD4 is a large membrane protein (717 aa) with two defined regions. The N-terminal region is predicted to have 12 transmembrane helices and might catalyze the polymerization of the TA precursors, the C-terminal LicD-type region adds at least one phosphorylcholine to the precursor chain repeating units (22,23). *S. oralis* Uo5 TacF shares only 47% sequence identity with TacF of *S. mitis*/*S. pneumoniae* (23). Our analysis revealed that four of the 138 strains do not contain the *S. mitis* but the *S. oralis* type IV LTA synthesis genes, consequently they are likely *S. oralis* strains (Table S1.1).

Interestingly, for 12/132 strains hits with >80% identity to TacF of both types were observed. 10 of 12 of these strains also contained the *S. mitis* RU-polymerase Smi_0770, in the two remaining strains neither the gene for Smi_0770 nor for LicD4 of *S. oralis* (Sor_0762) was present. For 42 of the 132 (31.8%) strains designated as *S. mitis* only a hit for *S. oralis* TacF was observed with high sequence identity (>80%). This suggests that *S. mitis* has more structural diversity in type IV TAs than has been previously anticipated.

The presence of two LTA molecules (types I and IV) in the analyzed *S. mitis* strains prompted us to investigate three aspects in detail. As the bioinformatics analysis indicated the presence of an LtaS-homolog of *S. aureus* in the *S. mitis* B6 (Smi_0753) genome (23), it is most likely that in *S. mitis* LtaS facilitates the synthesis of type I LTA. Therefore, we first searched for the corresponding *ltaS* gene in the genome of *S. mitis* NCTC10712 strain. Both proteins, Smi_0753 (of strain B6) and SMI10712_00646 (of strain NCTC10712), have 97% residues in common and are predicted to be polytopic membrane proteins with 5-transmembrane segments and a large C-terminal extracellular loop containing the complete sulfatase domain (PF00884), as does the LtaS protein (SAV0719) of *S. aureus* (24). Second, we used the *S. mitis* Smi_0753 protein sequence as query in tBlastn searches against the

*S. mitis* genomes available in NCBI microbial genome database (Table S1.1). Most *S. mitis* strains (100 out of 132, 75.8%) contain an LtaS homolog with 98% to 91% sequence identity. The 32 *S. mitis* genomes in which *ltaS* was absent contained the genes encoding proteins involved in the biosynthesis of *S. oralis*-like type IV TA, except three strains (Table S1.2). Third, we searched for LtaS homologs in related species, but we found no LtaS orthologs encoded by the *S. pneumoniae, S. oralis* and *S. infantis* genomes. This finding is in agreement with previous data (23,31), however 85 of 111 deposited *S. pseudopneumoniae* genomes (76.6%) contained an encoded *ltaS* gene (Table S2). For the remainder of 26 strains lacking the *ltaS* gene, 25 were not identified as *S. pseudopneumoniae* strains in the most recent phylogenetic analysis (32) (Table S2). All strains assigned as S. *pseudopneumoniae* contained the LtaS protein with a sequence identity of >97% to LtaS of *S. mitis* B6. Taken together, our genome analysis shows that LtaS homologs are present only in *S. mitis* and *S. pseudopneumoniae*. To further corroborate this, we selected representative strains from *S. pneumoniae, S. mitis and S. pseudopneumoniae* and analyzed the genomic localization of *ltaS*. We selected strains with available information about TA structures for four *S. pneumoniae* and three *S. mitis* strains (B6, NCTC10712, and SK137) as well as the type strains *S. mitis* NCTC12261 and *S. pseudopneumoniae* ATCC BAA-960. *S. pseudopneumoniae* IS7493 was selected due to the presence of a completed genome sequence. The *ltaS* gene is present as a monocistronic operon in *S. mitis* and *S. pseudopneumoniae* genomes (Fig. 7). All *S. pneumoniae* strains show a similar structure of this locus but lack the *ltaS* gene. The upstream and downstream regions are well conserved in all three species showing 90-93% identity at the nucleotide level. In addition, a BoxC element is located in the intergenic region between *spr1236* and *spr1237* genes in the *S. pneumoniae* genomes. Box elements are short repeated DNA sequences that are distributed randomly within the *S. pneumoniae* genomes (33). The function and origin of Box elements are still unknown, but they were found to enhance the genetic diversity and genome plasticity in *S. pneumoniae* (34).

**Figure 7.**
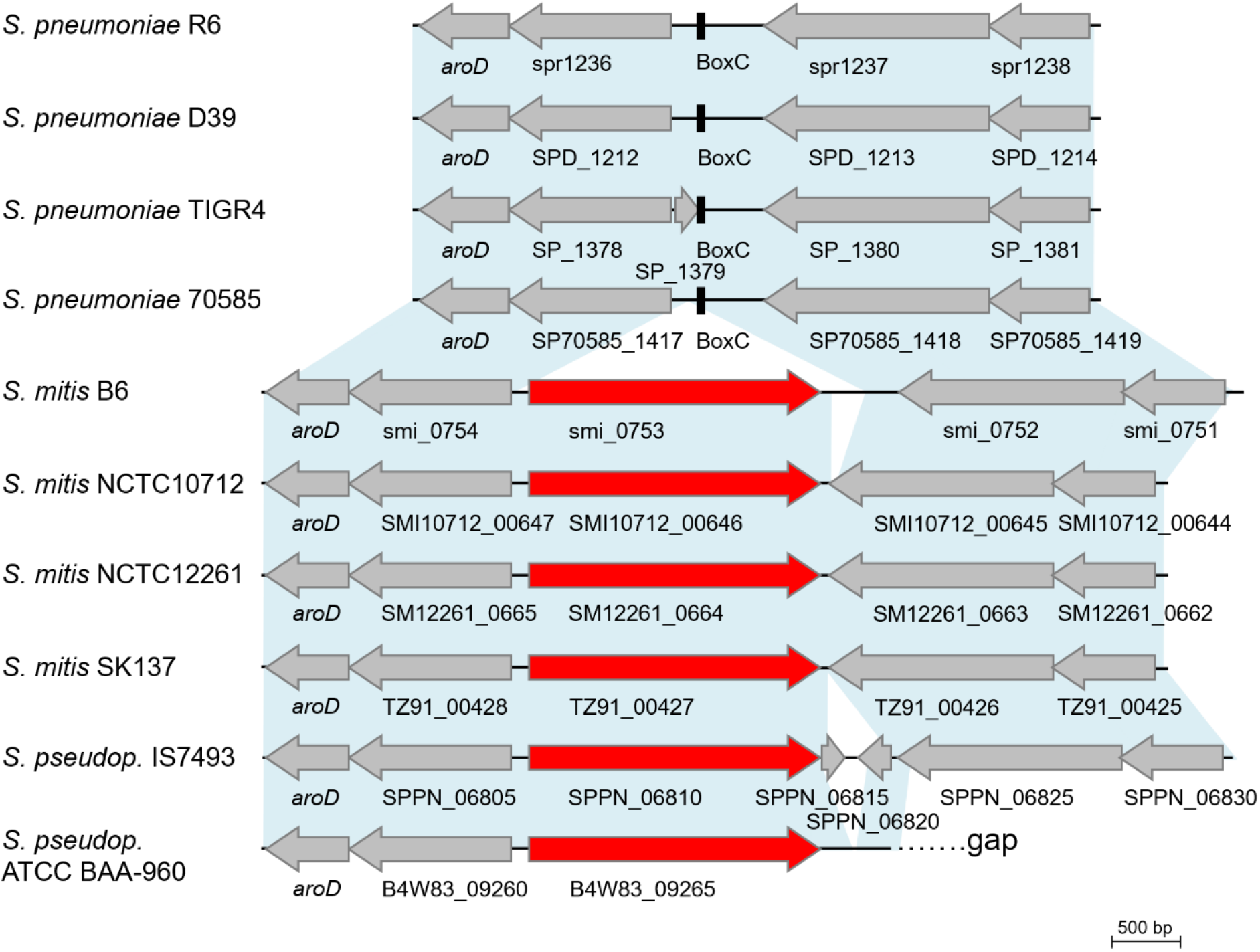
Comparison of the *ltaS* loci in three *Streptococcus* species. Representative strains of *S. mitis* and *S. pseudopneumoniae* carry an *ltaS* homolog (red). Upstream and downstream regions of the *ltaS* gene are conserved in all three species (genes are shown in grey and the old Gene/Locus nomenclature (sp/spr/smi) was used). Light blue areas illustrate BLASTn matches between sequences (90-93% identity). Large arrows show the gene orientation and are drawn in scale as indicated by the 500 bp bar. BoxC elements annotated in *S. pneumoniae* genomes are shown in black. The downstream region of the *ltaS* gene in the type strain of *S. pseudopneumoniae* ATCC BAA_960 is located in a gap. The NCBI accession numbers and references of strains used in this figure are listed in Table S3.

We also searched for putative promoter regions and putative transcriptional terminators of the *ltaS* gene in the B6 genome. Surprisingly, we found that the *ltaS* gene has two start codons, which are well conserved in all selected strains except *S. pneumoniae* (Fig. 8). It has been shown that in bacteria the predominant initiation codon is ATG (80.1%), while the alternative codon GTG is rarely occurring (11.6%) (35). However, for both start codons we distinctly predicted the promoter regions (Figure 8). The presence of two putative promoter regions and two putative translation start codons suggests a complex regulation of *ltaS* expression. The first promoter (start codon GTG) contains a -10 region, the putative ribosomal biding site, but the -35 element is not well conserved and is difficult to find. The second promoter (transcriptional start codon ATG) exhibits an extended -10 promoter element, which is four times more common in *S. pneumoniae* than in *E. coli* and can function without a -35 element (36). Remarkably, the four *S. pneumoniae* genomes contain the promoter region, although the *ltaS* gene is missing. Most likely, *S. pneumoniae* has lost the *ltaS* gene during evolution.

**Figure 8.**
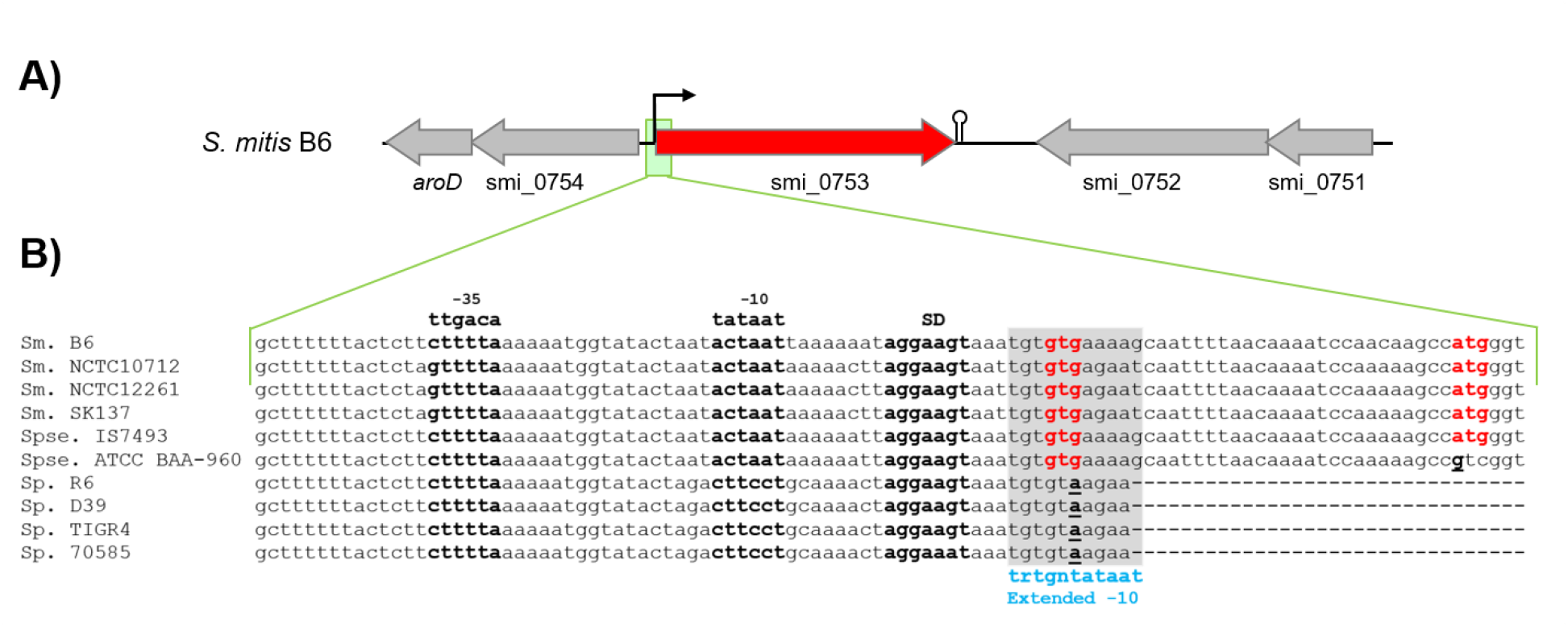
Putative promoter region of the *ltaS* in *S. mitis* strain B6. **A**) Genetic map of the *S. mitis* B6 genome region with *ltaS* (smi_0753). The *ltaS* gene is shown in red, the flanking genes in grey. The small black arrow indicates the putative promoter, the putative transcriptional terminator is shown as stem-loop structure. **B**) Comparison of the putative promoter region upstream of *ltaS* (smi_0753) with those of the other *S. pneumoniae* (Sp.), *S. mitis* (Sm.) and *S. pseudopneumoniae* (Spse.) strains. The alignment shows the conserved sequences located upstream of *ltaS* (smi_0753). The two putative start codons (GTG and ATG) are indicated in red. The putative promoter region (translation start site GTG) is highlighted in bold and the -10 and -35 elements as well as the putative ribosome-binding site (SD) are labelled. The extended -10 region (translation start ATG) is shown in bold within the grey area. The alterations within ATG start codon are underlined and bold. Sequences not present are represented in the alignment by (-).

## DISCUSSION

The present study provides the first detailed structural analysis of LTA molecules isolated from *S. mitis*. For our analysis, we have chosen the penicillin-sensitive strain NCTC10712 and the high-level beta-lactam resistant as well as multiple antibiotic resistant strain B6 as representatives. This structural investigation complements our earlier bioinformatics analysis of the TA-biosynthesis genes in members of the Mitis group (23), which suggested that the LTA of *S. oralis* should have an overall architecture similar to pneumococcal LTA with certain alterations due to a divergent enzymatic repertoire. We previously presented the *S. oralis* LTA structure and finally linked the observed structural variations between *S. oralis* and *S. pneumoniae* LTA to the distinct enzymes for TA biosynthesis (22). Based on these analyses (22,23) we predicted that *S. mitis* contains a type IV LTA with similar structure to that of *S. pneumoniae* (18), since LTA biosynthesis genes and their organization in the genomes of *S. pneumoniae* (R6) and *S. mitis* (B6) are almost identical (94-99% homology) with only one exception. The *smi1983* gene located in an operon equivalent to *spr0091-spr0092* encodes a glycosyltransferase that is similar to *S. pneumoniae* SP70585_0164 and *S. oralis* Sor_1862. The gene *sp70585_0164* was identified in the genome of the *S. pneumoniae* Sp70585 serotype 5 strain, which has been shown to incorporate galactose instead of glucose into its WTA (37) and Sor_1862 incorporates galactose into *S. oralis* LTA (22). The recently discovered lipoteichoic acid ligase TacL, which mediates the last step in LTA biosynthesis in *S. pneumoniae* (14) is conserved in both *S. mitis* strains with 97% homology. In summary, this suggested that *S. mitis* B6 contains a type IV LTA like pneumococci but with a galactose instead of the α-glucose. We confirmed this prediction by the structural analysis of the *S. mitis* LTA. Based on the BLAST search we predict that a subset of *S. mitis* strains with a homolog of Spr0091/SPD_0098 from *S. pneumoniae* R6/D39 contain exactly the same type IV LTA as *S. pneumoniae* (Table S1). This is consistent with results of a recent study (11), which showed that the majority of *S. mitis* strains possessed the equivalent of the *S. pneumoniae* 70585 gene SP70585_RS00795, as do all *S. oralis* and *S. infantis* strains, but it is exceptional amongst the *S. pneumoniae* strains. Moreover, two of the three identified glycosylglycerolipids of the *S. mitis* cell wall, α-glucopyranosyl-(1,3)-diacylglycerol and α-galactopyranosyl-(1,2)-α-glucopyranosyl-(1,3)-diacylglycerol, are identical to those present in *S. pneumoniae* (38). By contrast, the third identified glycosylglycerolipid, β-galactofuranosyl-(1,3)-diacylglycerol, is not present in *S. pneumoniae*. Even more remarkable, a second LTA polymer belonging to structure type I containing exactly this molecule as lipid anchor is present in both *S. mitis* strains investigated here. Besides *S. suis* (15), this is the second description of a Gram-positive bacterium being capable of synthezising two different LTA polymers. In the case of *S. mitis* the different LTAs are most likely synthezised via two distinct biosynthesis pathways, whereas in *S. suis* the synthesis of the two identified LTA types seems to be based on a single biosynthesis route. The type I LTA characterized in this study has been reported earlier in a *Streptococcus* sp. strain DSM 8747 (17), but the taxonomic characterization has never been published, thus the true genetic relatedness to *S. pneumoniae* remains unclear. Our finding of a type I LTA polymer in *S. mitis* strains contrasts an observation of Hogg *et al* (39) who suggested the presence of poly-glycerol-phosphate containing LTA in many oral streptococci, but not in *S. oralis* and *S. mitis* strains based on the detection of these molecules in phenol extracts of lyophilized cells using a poly-glycerol-phosphate-specific monoclonal antibody. A recent study (40) suggested the presence of a type I LTA besides the type IV LTA in *S. mitis* based on the identification of a potential biosynthetical intermediate, glycerophospho-diglycosyl-diacylglycerol (GPDGDAG). However, the authors of that study also reported the identification of GPDGDAG in *S. pneumoniae* and *S. oralis*, which do not contain type I LTA (18,22,41), what is further corroborated by the absence of LtaS in these species as elucidated here. Moreover, the mentioned work of Wei *et al* (40) showed by using an LtaS-knockout of *S. mitis* strain ATCC 49456 (SM61; NCTC12261) that GPDGDAG synthesis does not require LtaS. Notably, Wei *et al* could not identify, as earlier works (39), type I LTA by a poly-glycerol-phosphate-specific monoclonal antibody (40). We now clearly provide the missing evidence for a type I LTA in *S. mitis* by isolating this LTA and its subsequent structural analysis. Since the detected glycolipid anchor in the observed type I LTA is a mono-galactofuranosyl-DAG its synthesis will be independent from GPDGDAG synthesis, because this molecule contains a diglycosyl moiety and does not contain a galactofuranose. Our bioinformatics analysis aiming to identify the putative 1,2-diacylglycerol-3-β-galactofuranosyl transferase (or mono-galactofuranosyl-diacylglycerol synthase) involved in LTA glycolipid anchor biosynthesis in the *S. mitis* genomes did not provide a candidate enzyme. To the best of our knowledge such an enzyme has not been described in other bacteria so far.

Our bioinformatics analyses identified *ltaS* homologs only in *S. mitis* and *S. pseudopneumoniae* genomes, which distinguishes these from closely related species like *S. pneumoniae, S. oralis* and *S. infantis*. This is especially interesting regarding the human pathogen *S. pneumoniae*, which is genetically much closer related to *S. mitis* and *S. pseudopneumoniae* than *S. oralis* and *S. infantis*. The detailed and manual comparison of genomes done here is based on a limited number of genomes and requires as critical prerequisite the correct species assignment, which can be problematic, as seen by the recently reported misidentification of *S. pseudopneumoniae* (11,32). Here we selected specific LTA biosynthesis genes for an *S. mitis*/*S. pneumoniae* or an *S. oralis*-like LTA structure. At the current stage, this does not allow an unequivocal assignment of isolates to a specific species, which would require a higher number of curated genome sequences and the investigation of the associated TA structures. Therefore, future studies should focus on potentially new *S. mitis* subgroups, e.g., those containing homologs of both *S. mitis*- and *S. oralis*-type TacF, as well as the recently identified *S. oralis* subspecies. The higher structural diversity in TA biosynthesis within *S. mitis* has also been addressed recently by Kilian & Tettelin (11).

In summary, our work sets the stage for further investigations on the biosynthesis pathway of this particular type I LTA in *S. mitis*, the role of LtaS in its biosynthesis, and functional studies. Our work further suggests that a defined set of LTA biosynthesis genes could become a valuable metric to better differentiate between the closely related species studied here and more distant species such as *S. infantis* and the *S. oralis* subspecies *tigurinus* and *dentisanii*.

## EXPERIMENTAL PROCEDURES

### Bacterial strains and growth conditions

*S. mitis* B6 and NCTC10712 strains were grown at 37 °C without aeration in tryptone soya broth (TSB; Oxoid) medium and growth was followed by monitoring the absorbance at 600 nm. The strains were grown on D-agar plates supplemented with 3% defibrinated sheep blood (42). Cell wall preparation of *S. mitis* was performed as described previously (22). Briefly, bacteria were usually cultured in 5-liter batches of TSB medium harvested at late exponential phase by centrifugation (7,500 × g, 10 min) at room temperature. The cell pellet was washed with citric buffer (50 mM, pH 4.7) and resuspended in citric buffer containing 4% sodium dodecyl sulfate. The cell suspension was incubated for 20 min at 100 °C, stored at –80 °C and lyophilized subsequently.

### Extraction and isolation of LTA

LTA purification was performed as described elsewhere (14) twice independently for each strain. Yields of LTA preparations from 5 liters of bacterial culture were: for strain B6 4.9 mg and 12.1 mg; for strain NCTC10712: 4.9 mg (4 L culture) and 10.2 mg.

### Chemical treatments of LTA

Hydrazin treatment (to yield *O*-deacyl LTA) was performed following our earlier described procedure (15).

### Isolation of glycosylglycerolipids

For glycosylglycerolipids extraction, 126 mg dry bacterial pellet *S. mitis* strain B6 were suspended in 2 mL Millipore-water and 7.5 mL CHCl_3_/MeOH 1:2 v/v and thoroughly mixed (in a 50 mL Nalgene Oak Ridge Centrifuge Tube (FEP), Thermo Scientific). After 1 h shaking at RT 2.5 mL Millipore-water and 2.5 mL CHCl_3_ were added, mixed and shaken for another hour at RT. In parallel, for each of the four isolation batches 2 mL Millipore-water were treated in the same way to generate “equilibrated water”. The isolation mixtures were centrifuged at 4,000 × *g* for 10 minutes at 4 °C and the organic phases were transferred in fresh tubes. The remaining aqueous phase/interphase was extracted again with the organic phase of the “equilibrated water”-generation (10 s mixing, centrifugation as above). Resulting organic phases were combined with those of the first extractions. These combined phases were than washed with 4 mL of the “equilibrated water” (10 s mixing; centrifugation as above). The resulting organic phases were collected and dried under a stream of nitrogen (yielding 4.85 mg crude glycolipid extract). Glycosylglycerolipids were separated in two portions (applied as 5 µg/µL-solution in CHCl_3_/MeOH 2:1 v/v) by high-performance thin-layer chromatography (HPTLC) in CHCl_3_/MeOH 85:15 v/v on glass-backed 10 × 10 cm silica gel 60 F_254_ plates (Merck) and a small sidebar of the plate was stained with Hanessian’s stain (0.5 g of cerium(IV) sulphate tetrahydrate and 25 g of ammonium molybdate tetrahydrate in 471 ml of water supplemented with 29 ml of sulfuric acid with stirring (43)) at 150 °C. Appropriate bands were scrapped off from the silica (R_f_: 0.28-0.38, di-glycosyl-diacylglycerol; 0.56-0.65, mono-glycosyl-diacylglycerols) and recovered by extraction CHCl_3_/MeOH 2:1 v/v (30 s thorough mixing and subsequent centrifugation (1,080 × *g*, 10 min, 4 °C; three rounds of collecting and replacing the organic supernatant). After drying under a stream of nitrogen, the residual silica was removed by filtration in CHCl_3_/MeOH 2:1 v/v using a Acrodisc CR 13 mm Syringe Filter (0.2 µm PTFE membrane, PALL Life Sciences; washed with 2 × 2 mL CHCl_3_/MeOH 2:1 v/v prior to use).

### NMR spectroscopy

Deuterated solvents were purchased from Deutero GmbH (Kastellaun, Germany). NMR spectroscopic measurements were performed in D_2_O or deuterated 25 mM sodium phosphate buffer (pH 5.5; to suppress fast de-alanylation) at 300 K on a Bruker Avance^III^ 700 MHz (equipped with an inverse 5 mm quadruple-resonance Z-grad cryoprobe or with an inverse 1.7 mm triple-resonance Z-grad micro cryoprobe). Acetone was used as an external standard for calibration of ^1^H (δ_H_ = 2.225) and ^13^C (δ_C_ = 30.89) NMR spectra (44) and 85% of phosphoric acid was used as an external standard for calibration of ^31^P NMR spectra (δ_P_ = 0.00). Analysis of glycosylglycerolipids was performed in CD_3_OD and spectra were calibrated using the residual solvent peak (δ_H_ = 3.31, δ_C_ = 49.0) (44). All data were acquired and processed by using Bruker TOPSPIN V 3.1 or higher. ^1^H NMR assignments were confirmed by 2D ^1^H,^1^H-COSY and total correlation spectroscopy (TOCSY) experiments. ^13^C NMR assignments were indicated by 2D ^1^H,^13^C-HSQC, based on the ^1^H NMR assignments. Interresidue connectivity and further evidence for ^13^C assignment were obtained from 2D ^1^H,^13^C-heteronuclear multiple bond correlation and ^1^H,^13^C-HSQC-TOCSY. Connectivity of phosphate groups were assigned by 2D ^1^H,^31^P-HMQC and ^1^H,^31^P-HMQC-TOCSY.

### Mass spectrometry

All samples were measured on a Q Exactive Plus mass spectrometer (Thermo Scientific, Bremen, Germany) using a Triversa Nanomate (Advion, Ithaca, NY) as ion source. All measurements were performed in negative-ion mode using a spray voltage of -1.1 kV. Samples were dissolved in a water/propan-2-ol/trimethylamine/acetic acid mixture (50:50:0.06:0.02, v/v/v/v) in a final concentration of appr. 0.13 mg/mL. The mass spectrometer was externally calibrated with glycolipids of known structure. All mass spectra were charge deconvoluted and given mass values refer to the monoisotopic mass of the neutral molecules, if not indicated otherwise. Deconvoluted spectra were computed using Xtract module of Xcalibur 3.1. Software (Thermo, Bremen, Germany).

### Bioinformatic tools and analysis

The streptococcal genomic sequences used for the identification of genes involved in TA biosynthesis in the study are listed in Table S3 together with NCBI accession numbers and references. The accession numbers of the other *S. mitis* and *S. pseudopneumoniae* genomes used in this study are listed in Tables S1 and S2, respectively. In total 140 *S. mitis* and 112 *S. pseudopneumoniae* genomes were available in NCBI microbial genome database (as of 3rd January 2021). Two *S. mitis* strains (SK136 and NCTC12261) and *S. pseudopneumoniae* type strain ATCC BAA-960 (also known as CCUG 49455) were sequenced and deposited twice. Therefore, 138 different *S. mitis* and 111 *S. pseudopneumoniae* strains were included in the tBlastn analysis (45). The putative promoter regions were predicted by two consensus hexamers: -35 (5’
s-TTGACA-3’) and -10 (5’-TATAAT-3’) located upstream of the translation start codons of *ltaS*. Rho-independent transcriptional terminator was identified by Mfold program accessed through the mfold Web Server (46). Transmembrane helices and domain structure were predicted by using SMART Tool (http://smart.embl-heidelberg.de/) (47).

## Data availability

All data are contained within the manuscript.

## Acknowledgements

We gratefully acknowledge H. Käßner and B. Kunz (both RCB) for excellent technical assistance. We are indebted to N. Frankenberg-Dinkel for providing laboratory facilities to carry out the work of D.D. We would like to thank Reinhold Brückner for supplying the strains and helpful discussions. The work of D.D. was supported by the Deutsche Forschungsgemeinschaft (grant HA1011/11-3).

## ABBREVIATIONS

Ala: alanine
DAG: diacyl-glycerol
Gal: galactose
Glc: glucose
Gro: glycerol
HMQC: heteronuclear multiple quantum correlation
HSQC: heteronuclear single quantum correlation
LTA: lipoteichoic acid
Ppm: parts per million
RU: repeating unit
TOCSY: total correlation spectroscopy.

