## Supplemental Figures 1 and 2, Supplemental Table 3 for "Commensal *Streptococcus mitis* produces two different lipoteichoic acids of type I and type IV"

**Materials included**
Figure S1
Figure S2
Table S1 (separate file)
Table S2 (separate file)
Table S3

### Figure S1

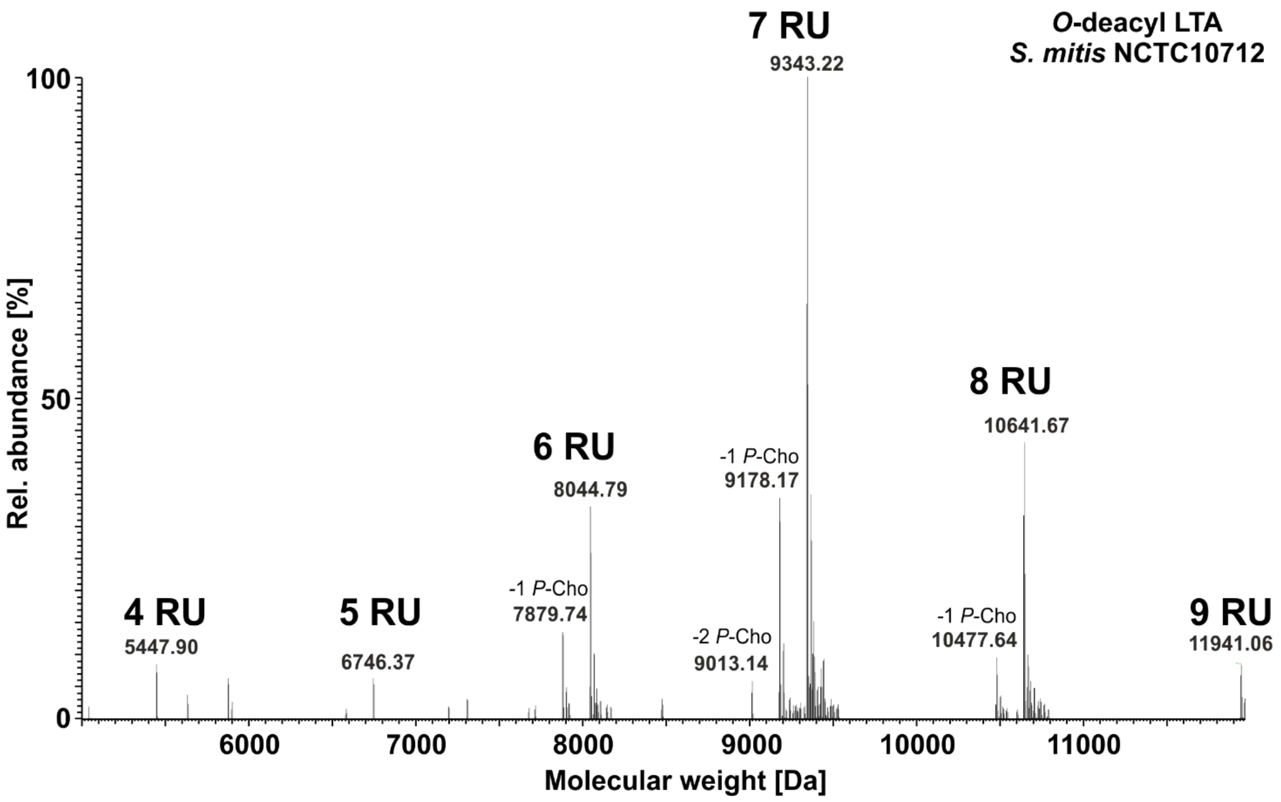

**Figure S1. MS analysis of one batch of *O*-deacyl LTA isolated from *S. mitis* strain NCTC10712.** Section of the charge deconvoluted MS spectrum (acquired in negative-ion mode) typical for type IV LTA is shown.

### Figure S2

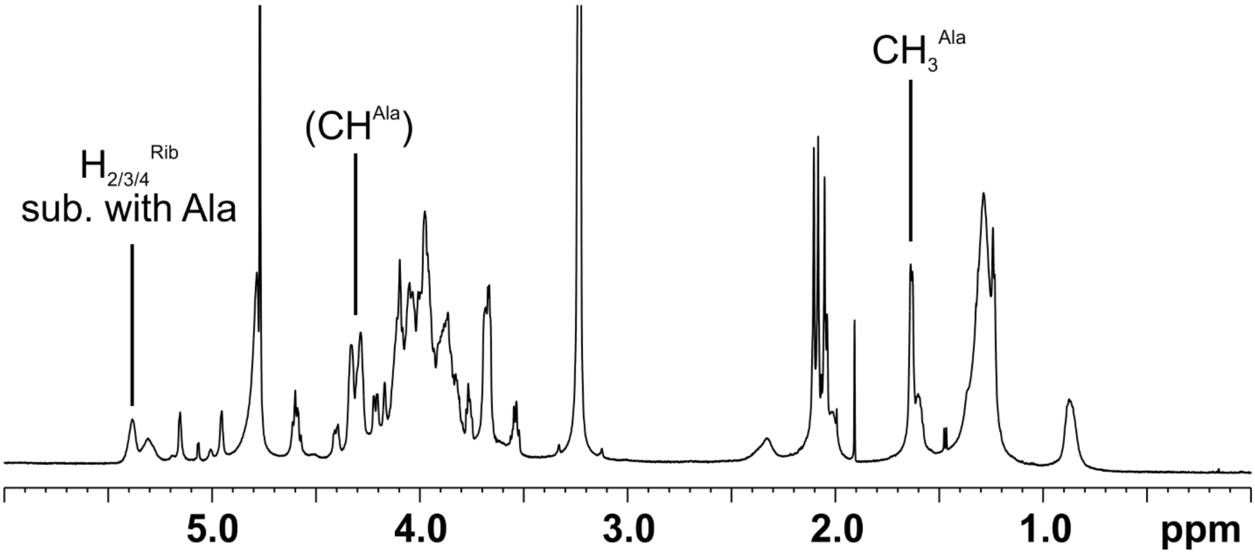

**Figure S2. [1]H NMR analysis of native LTA of *S. mitis* strain NCTC10712 reveals alanylation of LTA.** Shown is the $^1$H NMR spectra ($\delta_H$ 6.0-0.0) of LTA isolated from *S. mitis* strain NCTC10712 recorded in deuterated 25 mM sodium phosphate buffer (pH 5.5) at 300 K. The signals originating from the methyl ($CH_3$) group of alanine at $\delta_H$ 1.63 ppm and from protons H-2, H-3 and/or H-4 (specific position is unknown) of alanylated ribitol at $\delta_H$ 5.43-5.35 ppm are clearly indicative for alanylated LTA. The CH-group of alanine at $\delta_H$ 4.32-4.26 ppm is overlaid mainly by signals originating from $CH_2O$ of *P*-Cho substituents.

### Table S3

**Table S3. Genomes of *Streptococcus* strains used for bioinformatics analysis in this study.**

| Species | Strain | Genome size (mb) | No. of contigs | Accession | Reference |
|---|---|---|---|---|---|
| *S. pneumoniae* | D39 | 2,05 | 1 | CP000410 | (1) |
| | R6 | 2,04 | 1 | AE007317 | (2) |
| | TIGR4 (ATCC BAA-334) | 2,16 | 1 | AE005672 | (3) |
| | 70585 | 2,18 | 1 | NC_012468.1 | (4) |
| *S. mitis* | B6 | 2,15 | 1 | NC_013853 | (5) |
| | NCTC10712 | 1,84 | 44 | LROT00000000.1 | (6) |
| | NCTC12261/SK142$^{T}$/ATCC49456 | 1,87 | 1 | CP028414 | (7) |
| | SK137 (CCUG 35791) | 1,98 | 41 | JPFS00000000 | (8) |
| *S. pseudo-pneumoniae* | IS7493 | 2,20 | 2 | CP002925 | (9) |
| | ATCC BAA-960$^{T}$ | 2,09 | 86 | NZ_MWSM00000000.1 | * |

$^{T}$ Type strain; (*) - Direct submission in NCBI
